# The effect of combined patching and citalopram on visual acuity in adults with amblyopia: a randomized, crossover, placebo-controlled trial

**DOI:** 10.1101/587980

**Authors:** Alice K. Lagas, Joanna M. Black, Bruce R. Russell, Robert R. Kydd, Benjamin Thompson

## Abstract

Non-human animal models have demonstrated that selective serotonin reuptake inhibitors (SSRIs) can enhance plasticity within the mature visual cortex and enable recovery from amblyopia. The aim of this study was to test the hypothesis that the SSRI citalopram combined with part-time patching of the fellow fixing eye would improve amblyopic eye visual acuity in adult humans. Following a cross-over, randomized, double blind, placebo-controlled design (pre-registration: ACTRN12611000669998), participants completed two 2-week blocks of fellow fixing eye patching. One block combined patching with citalopram (20 mg/day) and the other with a placebo tablet. The blocks were separated by a 2-week washout period. The primary outcome was change in amblyopic eye visual acuity. Secondary outcomes included stereoacuity and electrophysiological measures of retinal and cortical function. Seven participants were randomized, fewer than our pre-specified sample size of 20. There were no statistically significant differences in amblyopic eye visual acuity change between the active (mean ± SD change = 0.08±0.16 logMAR) and the placebo (mean change = −0.01±0.03 logMAR) blocks. No treatment effects were observed for any secondary outcomes. However, 3 of 7 participants experienced a 0.1 logMAR or greater improvement in amblyopic eye visual acuity in the active but not the placebo block. These results from a small sample suggest that larger-scale trials of SSRI treatment for adult amblyopia may be warranted. Considerations for future trials include drug dose, treatment duration and recruitment challenges.

## Introduction

Disruptions to binocular vision such as strabismus (an eye turn) or anisometropia (unequal refractive error between the two eyes) during the critical period of visual development can cause a neurodevelopmental disorder of vision called amblyopia [1, 2]. The deficits associated with amblyopia encompass a wide range of monocular and binocular visual functions [3, 4] and also extend to the fellow fixing eye [5]. Clinically, amblyopia is typically diagnosed on the basis of a monocular visual acuity loss that cannot be explained by ocular pathology combined with an amblyogenic factor [1]. Current treatments for amblyopia in childhood involve the provision of refractive correction followed by patching or penalization of the fellow fixing eye to promote use of the amblyopic eye. These treatments are effective [6–12], but efficacy appears to decline with increasing age in children [13–16], possibly due to a decline in neural plasticity as the visual cortex matures and exits the critical period for visual development [17–20]. A growing body of literature demonstrates that vision can improve in adult humans with amblyopia through interventions such as monocular [21, 22] and binocular [23–27] perceptual learning and non-invasive brain stimulation [28–32]. However, these approaches have not yet translated into positive randomized clinical trials in adult patients that are required for translation into clinical practice [33].

Amblyopia also forms the basis of a prominent non-human animal model for studying cortical development and plasticity [34]. Monocular amblyopia can be induced in non-human animals within the critical period of visual development using an eyelid suture, induction of strabismus, or provision of anisometropic refractive error [35]. Over the past decade or so, a considerable number of studies have used this model to explore post-critical period neuroplasticity [36]. Successful interventions for amblyopia recovery in post-critical period animal models include dark exposure [37, 38], enriched visual environments [39], food restriction [40], binocular training [41], physical exercise [42] and retinal inactivation [43].

Pharmaceutical interventions have also been investigated in rodent models of amblyopia. A particularly striking result was reported by Maya Vettencourt et al. [44] whereby chronic administration of the selective serotonin reuptake inhibitor (SSRI) fluoxetine enabled recovery of normal visual cortex responses and visual acuity in mature rats with unilateral deprivation amblyopia. This effect occurred when fluoxetine was administered before and during eyelid suture of the non-deprived eye and opening of the deprived eye (a procedure known as a reverse suture). The improvements in visual function were linked to reduced GABA mediated inhibition within the visual cortex and increased expression of brain derived neurotrophic factor (BDNF). This finding is of particular interest in the context of amblyopia treatment in adult humans because SSRIs are widely available to clinicians. Furthermore, SSRIs may enhance plasticity within the human motor [45, 46] and visual [47] cortex. Fluoxetine has also been found to enhance physiotherapy outcomes after stroke, possibly by increasing cortical plasticity [48]. However, fluoxetine did not enhance visual perceptual learning of a motion discrimination task or motor cortex plasticity in a study of healthy human adults [49].

Two studies have investigated the use of fluoxetine to treat human amblyopia. Sharif et al. [50] compared 3 months of fellow fixing eye patching plus fluoxetine (0.5mg/kg/day, n = 20) to patching plus a placebo tablet placebo (n = 15) in older children and adults (10-40 years) with amblyopia. A significantly greater amblyopic eye visual acuity improvement in the fluoxetine compared to the placebo group was observed. However, Huttunen et al. [51] found no differences in visual function improvement between a group of adults with amblyopia treated for 10 days with combined perceptual learning and fluoxetine (20 mg per day, n = 22) and a group treated with perceptual learning combined with a placebo tablet (n = 20).

In this study we explored the effects of 2 weeks (14 days) of the SSRI citalopram combined with fellow fixing eye patching on visual acuity, stereopsis and visually evoked retinal and cortical responses in adults with amblyopia. We anticipated that recruitment would be challenging due to the use of patching and the administration of an anti-depressant drug. We therefore adopted a placebo-controlled, randomized, double blind, crossover design. In this context citalopram was chosen over fluoxetine (as used in prior studies) because citalopram has a shorter half-life [52] that allowed for a manageable washout period to be incorporated into the design of the study. No significant effects of citalopram were observed, although our study may have been underpowered due to recruitment challenges.

## Methods

### Trial design

The single-site trial involved two blocks of fellow fixing eye patching each lasting two weeks separated by a two-week washout period. Participants were provided with citalopram (1 x 20mg tablet per day) during one patching block and otherwise identical placebo tablets (sucrose) during the other block. Block order was randomized using a random number generator. The timing of baseline and outcome measures are shown in Figure 1. Only the pharmacist dispensing the tablets, who did not interact with study participants, was unmasked to block order. Study participants and all other members of the research team were masked to the randomization. The study was approved by the Northern X Regional Ethics Committee in New Zealand (NTX/11/06/044) and pre-registered as a clinical trial (ACTRN12611000669998).

**Figure 1.**
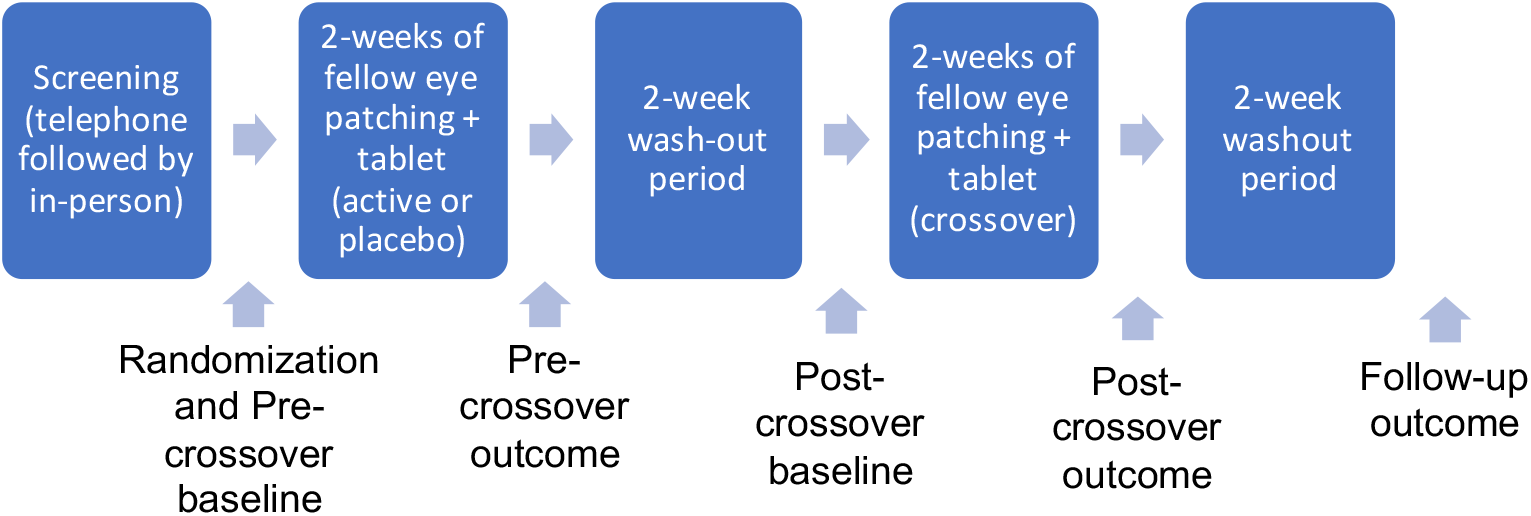
Schematic of the study protocol and the timing of baseline, outcome and follow-up measures.

Participants completed a screening protocol consisting of a telephone interview followed by a full optometric examination, medical history, the Profile of Mood State Short Form questionnaire (POMS-SF) and the Depression Anxiety and Stress Scale (DASS-21).

Study inclusion criteria were: 18 years of age or over, 0.2 logMAR or worse visual acuity in the amblyopic eye, 0.0 logMAR or better visual acuity in the fellow fixing eye, an interocular acuity difference of at least 0.2 logMAR, the presence of a strabismus and/or anisometropia defined as a difference in spherical equivalent refractive error of 1.5 dioptres or greater between the eyes. The exclusion criteria were: the presence of ocular pathology, an explanation for the visual acuity loss other than amblyopia, personal or family history of a mood disorder, diabetes, history of addiction, current use of medications or supplements known to alter mood, medications that interact with SSRIs such as codeine and abnormal mood states evident on the mood questionnaires as reviewed by a psychiatrist. Prior to randomization, participants who were not wearing optimal full correction for both eyes were provided with full correction (either spectacles or contact lenses) and were reviewed every four weeks until visual acuity was stable (<0.2 logMAR difference between visits). Participants were recruited through the University of Auckland Optometry Clinic, referral from eye care practitioners, word of mouth and newspaper advertisements. Participants were compensated for their time.

### Baseline and outcome measures

Visual acuity (VA) was assessed using a computerized ETDRS chart (Medmont) from 6 m. The right eye was tested first. Each correctly identified letter was worth 0.02 logMAR. Binocular vision was assessed using a unilateral cover test, a prism cover test, the Worth 4-dot test (33 cm and 6 m) and the TNO stereoacuity test. Electrophysiological measurements of retinal and visual cortex function were made using ISCEV-standardized electrophysiological protocols on a Roland Retiscan system (software version 4.13.1.8). The following tests were applied monocularly (right eye first); pattern-ERG (1° check size – modified from the 0.8° standard for direct comparison with the VEP stimuli), VEP (1° and 0.3° check sizes), multifocal ERG with pupil dilation. ERG measures were included so that any retinal effects of citalopram could be accounted for if the trial was positive. The POMS-SF questionnaire was completed at each study visit and participants completed a patching diary for each 2-week patching session. Brain derived neurotropic factor (BNDF) phenotype has been identified as a possible mediator of cortical plasticity [53] and BNDF upregulation has been identified as a mechanism for increased visual cortex plasticity following fluoxetine administration in rats [44]. To test for BDNF polymorphisms, participants provided a blood sample directly after the first two-week block of patching. Following a previously reported protocol [49], an Agena MassArray IPLEX assay (Agena Bioscience, San Diego, CA, USA) was used for genotyping. A Brucker Mass Spectrometer with optimized parameters for iPLE× chemistry was then used to resolve single base extensions. Typer 4 analysis software (Agena Bioscience) enabled visual inspection of generated peaks in comparison to the non-template control.

### Statistical analysis

At the time of study initiation, no previous studies of SSRIs in human amblyopia treatment were available. Therefore, we selected a sample size of 20 based on recruitment estimates for the study site. Outcome measures were analysed separately using mixed ANOVAs with within-subject factors of Session (baseline vs. outcome) and Treatment (active vs. placebo) and a between subjects factor of Group (active first vs. placebo first). Data Availability: all clinical data are provided within the manuscript tables.

## Results

Sixty-one participants expressed interest in the study and were sent a study information package. Twenty-eight participants responded and were assessed for eligibility. Seven participants were randomized. The CONSORT diagram for these participants is shown in Figure 2. Reasons for exclusion included time commitment too great, medical or recreational use of drugs, vision too good in the amblyopic eye and diabetes. One participant who did not meet the visual acuity inclusion criteria was randomized (P6, see Table 1). Data from this participant were included in the final analysis due to the small sample size. Randomized participant details, including BDNF polymorphism, are shown in Table 1.

**Figure 2.**
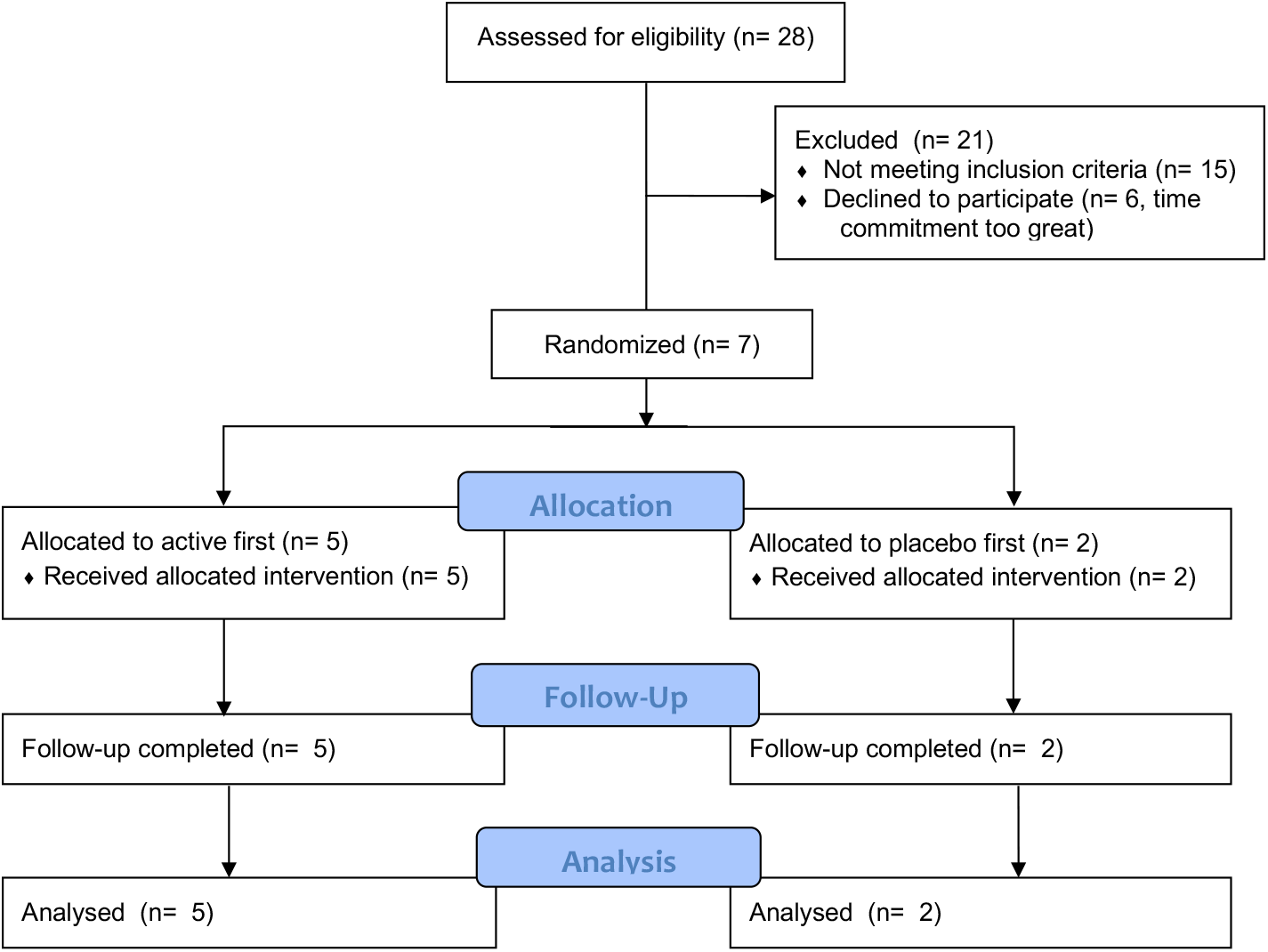
CONSORT diagram for the study.

**Table 1.**
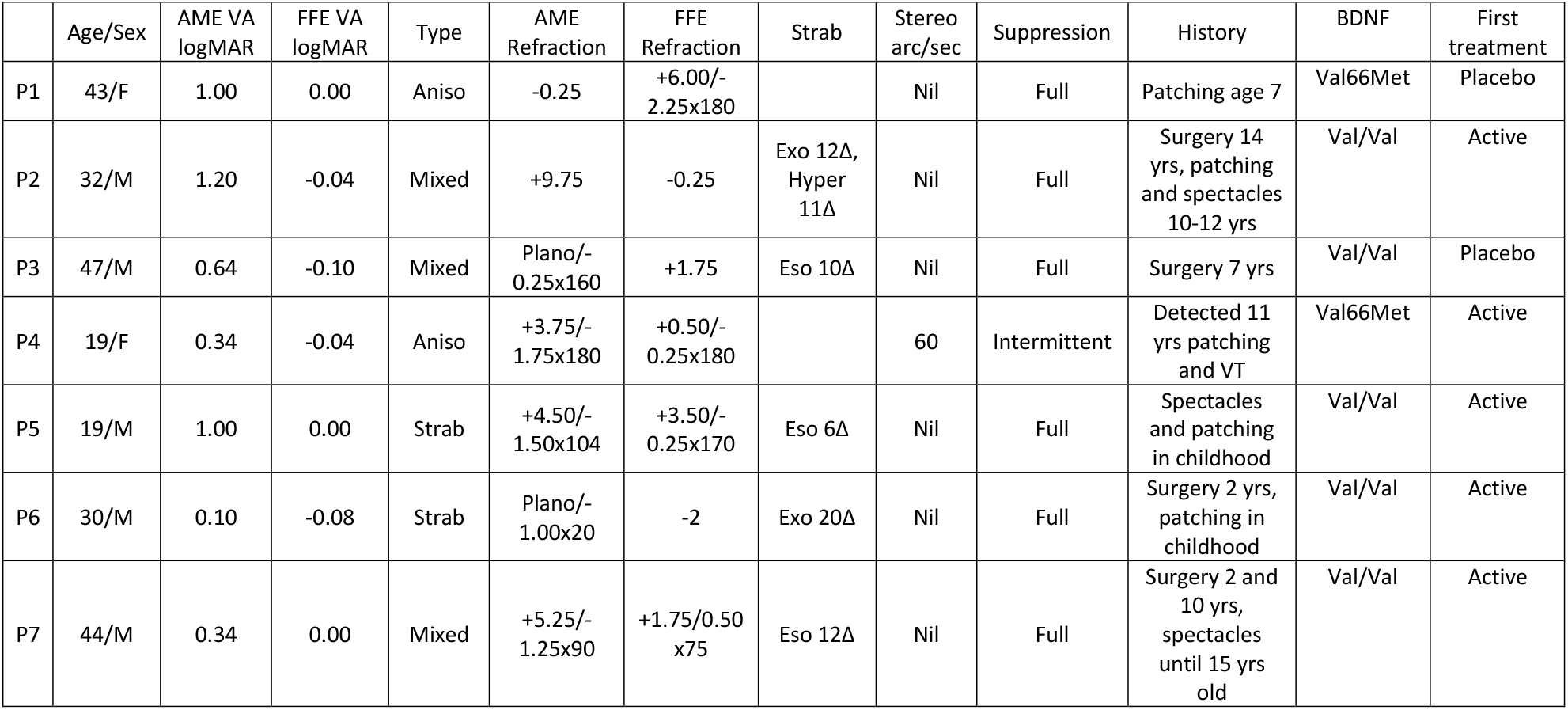
Participant details. AME, amblyopic eye; FFE, fellow fixing eye; VA, visual acuity; exo, exotropia; eso, esotropia; hyper, hypertropia; aniso, anisometropic; strab, strabismic/strabismus; VT, vision training; BDNF, brain derived neurotrophic factor.

|  | Age/Sex | AME VA<br>logMAR | FFE VA<br>logMAR | Type | AME<br>Refraction | FFE<br>Refraction | Strab | Stereo<br>arc/sec | Suppression | History | BDNF | First<br>treatment |
| --- | --- | --- | --- | --- | --- | --- | --- | --- | --- | --- | --- | --- |
| P1 | 43/F | 1.00 | 0.00 | Aniso | -0.25 | +6.00/-<br>2.25x180 |  | Nil | Full | Patching age 7 | Val66Met | Placebo |
| P2 | 32/M | 1.20 | -0.04 | Mixed | +9.75 | -0.25 | Exo 12Δ,<br>Hyper<br>11Δ | Nil | Full | Surgery 14<br>yrs, patching<br>and spectacles<br>10-12 yrs | Val/Val | Active |
| P3 | 47/M | 0.64 | -0.10 | Mixed | Plano/-<br>0.25x160 | +1.75 | Eso 10Δ | Nil | Full | Surgery 7 yrs | Val/Val | Placebo |
| P4 | 19/F | 0.34 | -0.04 | Aniso | +3.75/-<br>1.75x180 | +0.50/-<br>0.25x180 |  | 60 | Intermittent | Detected 11<br>yrs patching<br>and VT | Val66Met | Active |
| P5 | 19/M | 1.00 | 0.00 | Strab | +4.50/-<br>1.50x104 | +3.50/-<br>0.25x170 | Eso 6Δ | Nil | Full | Spectacles<br>and patching<br>in childhood | Val/Val | Active |
| P6 | 30/M | 0.10 | -0.08 | Strab | Plano/-<br>1.00x20 | -2 | Exo 20Δ | Nil | Full | Surgery 2 yrs,<br>patching in<br>childhood | Val/Val | Active |
| P7 | 44/M | 0.34 | 0.00 | Mixed | +5.25/-<br>1.25x90 | +1.75/0.50<br>x75 | Eso 12Δ | Nil | Full | Surgery 2 and<br>10 yrs,<br>spectacles<br>until 15 yrs<br>old | Val/Val | Active |

Baseline and outcome data for amblyopic eye visual acuity are shown in Table 2. There was no significant interaction between Session and Treatment (F_1,5_ = 1.7, p = 0.25, partial η^2^ = 0.26) indicating no difference between the active and placebo treatment. There was also no main effect of Session indicating the absence of a visual acuity improvement across the two periods of patching (F_1,5_ = 1.7, p = 0.25, partial η^2^ = 0.26). Overall, no main effects or interactions were significant in the analysis (all p > 0.25). An inspection of individual data (Table 2) indicated that 3/7 participants improved by >0.1 logMAR line in the active but not placebo condition. One of these participants had a val66met BDNF polymorphism. The remaining two had val/val BNDF polymorphisms. No main effects or interactions were present for the fellow fixing eye visual acuity data (all F < 3.9, all p > 0.1, all partial η^2^ < 0.4).

**Table 2.** Amblyopic eye visual acuity results. Change values were calculated by subtracting outcome from baseline. All values are in logMAR.

|  | Active<br>Baseline | Active<br>Outcome | Active<br>Change | Placebo<br>Baseline | Placebo<br>Outcome | Placebo<br>Change | Final<br>Washout |
| --- | --- | --- | --- | --- | --- | --- | --- |
| P1 | 0.94 | 0.82 | 0.12 | 1 | 0.97 | 0.03 | 0.87 |
| P2 | 1.20 | 1.20 | 0.00 | 1 | 1 | 0 | 1.1 |
| P3 | 0.73 | 0.60 | 0.13 | 0.64 | 0.67 | -0.03 | 0.633 |
| P4 | 0.34 | 0.36 | -0.02 | 0.32 | 0.32 | 0 | 0.3 |
| P5 | 1.00 | 0.60 | 0.40 | 0.866 | 0.9 | -0.034 | 0.74 |
| P6 | 0.10 | 0.10 | 0.00 | 0.1 | 0.08 | 0.02 | 0.14 |
| P7 | 0.34 | 0.40 | -0.06 | 0.32 | 0.36 | -0.04 | 0.32 |
| **Mean**<br>**(SD)** | **0.66**<br>**(0.36)** | **0.58**<br>**(0.35)** | **0.08**<br>**(0.16)** | **0.61**<br>**(0.36)** | **0.61**<br>**(0.36)** | **-0.01**<br>**(0.03)** | **0.59**<br>**(0.35)** |

Adherence data are shown in Table 3. Adherence did not differ significantly between active and placebo blocks (t_6_ = 1.0, p = 0.9). On average participants had approximately 70% adherence with the 120 minutes per day of prescribed patching. There was no correlation between patching adherence and visual acuity change in either the active (r_7_ = −0.2, p = 0.6) or placebo blocks (r_7_ = 0.3, p = 0.5).

**Table 3.** Self-reported patching adherence data sourced from participants’ patching diaries. Data are shown as mean minutes of patching per day (SD). The prescribed dose was 120 minutes per day.

|  | Active | Placebo | Difference |
| --- | --- | --- | --- |
| P1 | 116<br>(31) | 114<br>(24) | 2 |
| P2 | 40<br>(19) | 29<br>(21) | 11 |
| P3 | 55<br>(28) | 55<br>(7) | 1 |
| P4 | 96<br>(38) | 111<br>(32) | -15 |
| P5 | 75<br>(0) | 75<br>(0) | 0 |
| P6 | 111<br>(78) | 111<br>(74) | 0 |
| P7 | 111<br>(32) | 111<br>(32) | 0 |
| **Mean**<br>**(SD)** | **86**<br>**(30)** | **87**<br>**(34)** | **0.1**<br>**(7.6)** |

Only participant P6 exhibited a change in stereoacuity, improving from nil to 240 arc /sec in the active block and from nil to 480 arc/sec in the placebo block. Follow-up stereoacuity was nil. No significant treatment effects were evident for any of the electrophysiological measurements (all F < 2.0, all p > 2). Figure 3 shows 1° check stimulus VEP latencies (left) and N75-P100 amplitudes (right) for both the amblyopic and fellow fixing eyes. Figure 4 shows example multifocal ERG data for participant P7 (first baseline measure) and figure 5 shows example pattern ERG and VEP data for the same participant. There were no treatment effects on POM-SF scores.

**Figure 3.**
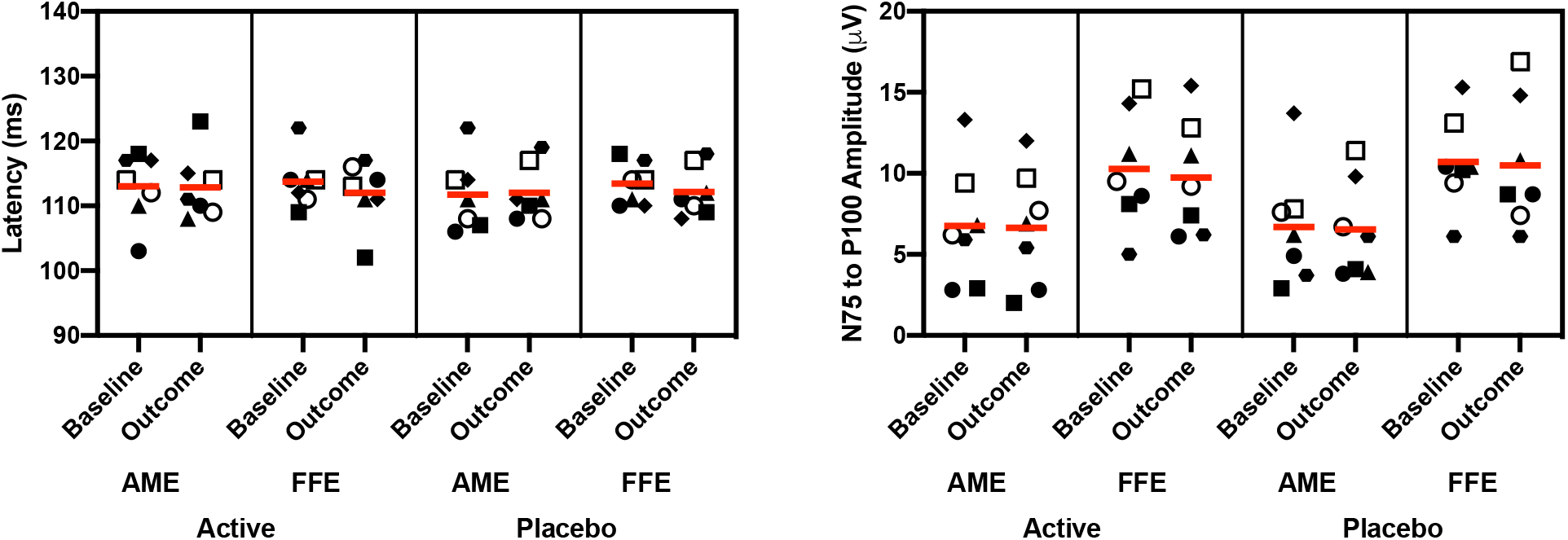
VEP results for the 1° check stimulus. Latencies for the P100 component are shown on the left and amplitudes for the N57-P100 waveform component are on the right. Individual participants are shown with different symbols: P1-7 as follows; filled circle, filled square, filled triangle, filled diamond, filled hexagon, open circle, open square. Horizontal lines depict group mean values.

**Figure 4.**
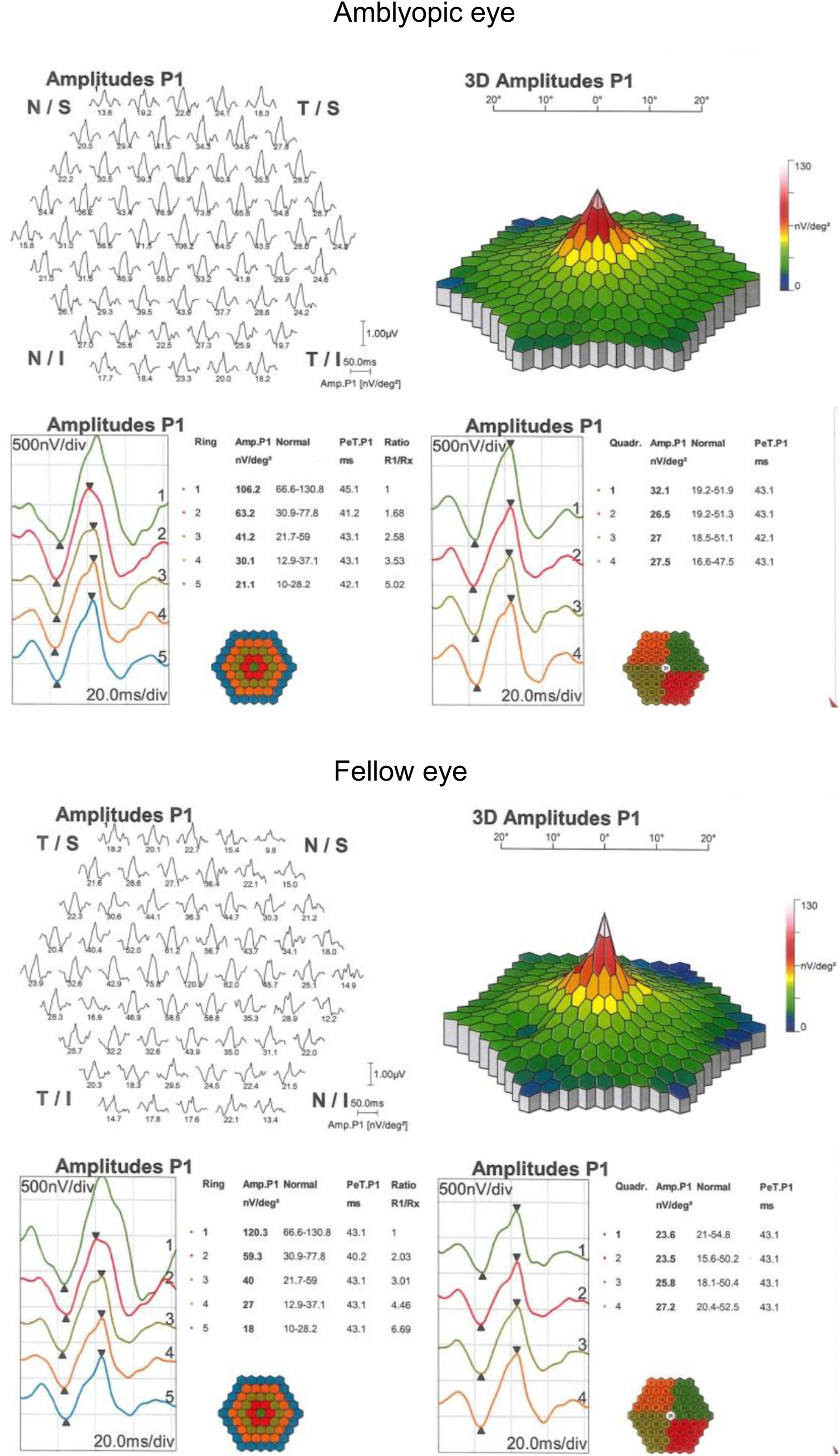
Multifocal ERG results for participant P7 (first baseline session).

**Figure 5.**
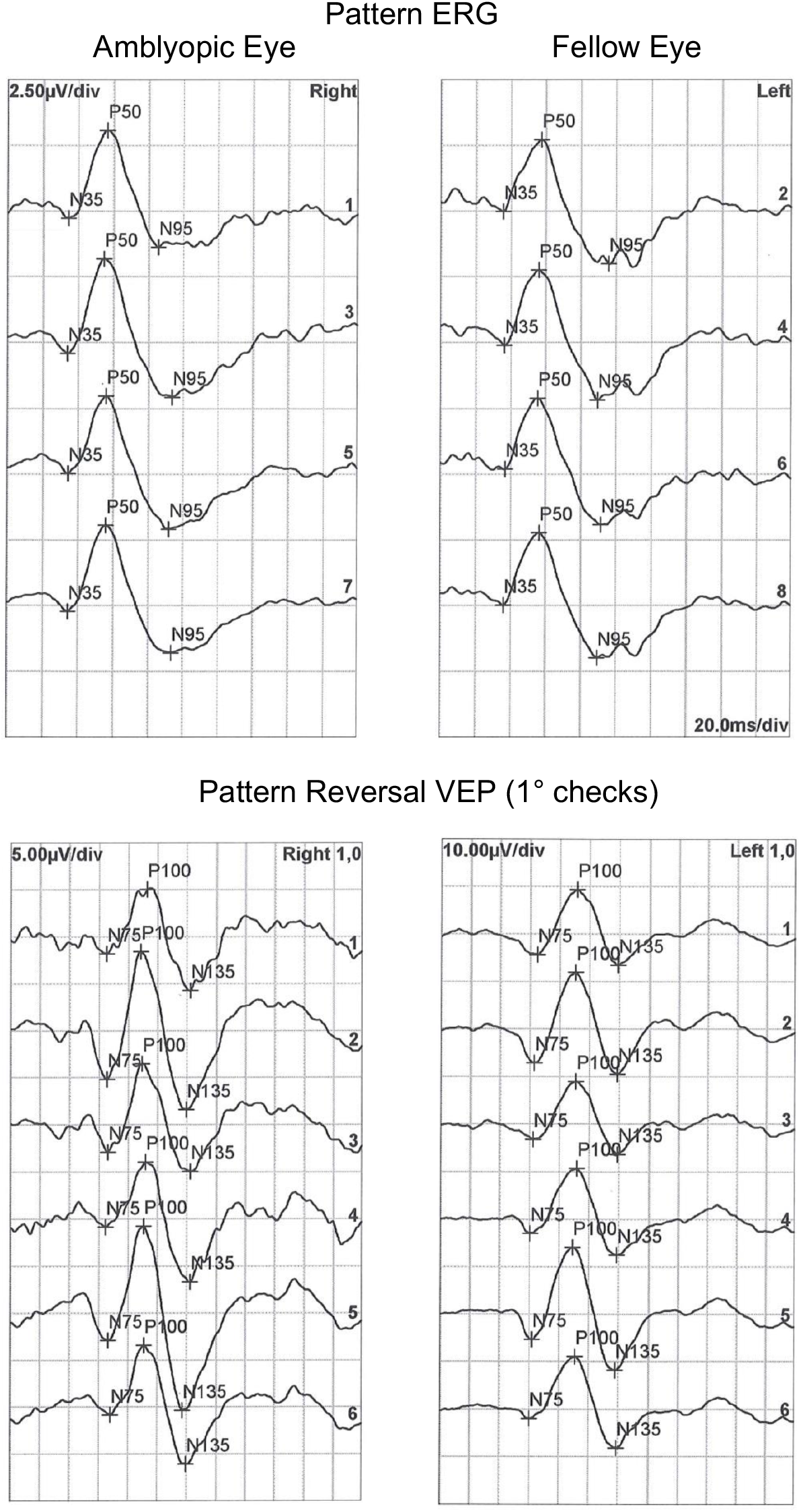
Pattern ERG (top) and pattern reversal VEP (bottom) results for the amblyopic (left) and fellow (right) eyes of participant P7 (first baseline session).

## Discussion

The SSRI fluoxetine enabled recovery of vision in mature rats with amblyopia [44] and has recently been reported to enhance the effect of patching in older children and adults [50]. We found no effect of the SSRI citalopram combined with two weeks of patching on amblyopic eye visual acuity or a range of secondary outcome measures in adults with amblyopia. These results are broadly consistent with another recent study with a similar duration treatment period (10 days) that reported no advantage of combining fluoxetine with perceptual learning compared to perceptual learning alone in adults with amblyopia [51]. A preliminary study of donepezil [54] and a randomized clinical trial of levodopa [55] have also found no benefit of drug treatment in amblyopia therapy. In addition, we found no effect of two weeks of patching alone in adult patients despite reasonable self-reported adherence. This is expected based on the short treatment period and the reduced effect of patching with increasing age [15, 16, 56].

A number of factors may explain the lack of a drug treatment effect in our study. First, and perhaps most importantly, we did not achieve our planned sample size of 20 participants due to difficulties with recruitment. This led to a small sample with varied amblyopia etiology and treatment history. Barriers to recruitment included the time commitment required by the study and the stringent medical inclusion criteria. Therefore, our study may be underpowered to detect a treatment effect, although the sample size is within the range of previous case-series perceptual learning studies that have reported treatment effects [57]. It is intriguing that three participants exhibited an amblyopic eye visual acuity improvement of 0.1 logMAR or greater for the active but not the placebo treatment sessions. These participants had relatively poor baseline amblyopic eye visual acuity compared to most of the other participants. No participants exhibited any improvement for the placebo sessions. This observation suggests that further testing of SSRI treatment effects in adults with amblyopic may be warranted.

Whereas previous studies have used fluoxetine, we used citalopram because it has a short lead in period of two hours [58]. Moreover, citalopram has a shorter half-life than fluoxetine; the distribution phase lasts about ten hours and the terminal half-life (T1/2) is 30-35 hours for citalopram [58] in contrast to two to four days half-life for fluoxetine [59]. Citalopram and fluoxetine appear to have the same efficacy for treating major depression [60] and comparable effects on plasma GABA, glutamine and glutamate levels in human patients [61]. However, citalopram and fluoxetine have different patterns of binding affinity within the human brain [62]. It is currently unknown whether the two drugs differ in the extent to which they promote visual cortex plasticity.

We used a 20 mg/day dose of citalopram over 2 weeks. It is possible that larger doses and longer treatment times are required to replicate the effects found in non-human animals. Supporting this idea, Sharif et al. [50] found a significant effect of combined fluoxetine and patching with dose of 0.5 mg/kg/day and a 3-month treatment period whereas Huttunen et al [51] found no effect with 20 mg per day over 10 days. The parameter space for dosing and treatment duration is large for drug intervention studies of this type and further work is required to identify optimal values. In addition, genotype may also influence an individual’s response to a pharmacological intervention. In this study we measured BDNF polymorphisms because they have been linked to neuroplasticity [53] and an increase in BDNF expression has been identified as a key mechanism in SSRI-induced recovery from amblyopia in mature rats [44]. There was no relationship between BDNF polymorphism and treatment response in this study with both val/val and val66met carriers improving by 1 logMAR line or more. However, the small sample size precludes any strong conclusions.

In agreement with Huttunen et al. [51], we found no effect of SSRI treatment on VEP parameters. This is in contrast to other emerging potential approaches to amblyopia treatment in adulthood such as the non-invasive brain stimulation technique anodal transcranial direct current stimulation that increases VEP amplitude [28]. The lack of any VEP changes is consistent with the lack of a treatment effect on any of the other outcome measures used within this study. Retinal electrophysiology was also conducted to rule out any retinal changes if a positive treatment effect was observed. No retinal changes were observed, in agreement with the overall study results.

In addition to the small sample size, a weakness of our study is that one participant (P6) did not meet the visual acuity inclusion criteria. We retained this participant in the study due to difficulties with recruitment. We note that excluding this participant from the sample does not change the pattern of results.

In conclusion, we found no effect of 2 weeks of combined citalopram and patching on amblyopic eye visual acuity in adults with amblyopia. This result may have been due to our study being underpowered as a result of recruitment challenges. Three out of seven participants did exhibit an amblyopic eye visual acuity improvement of 0.1 logMAR or more with combined citalopram and patching suggesting that further studies in this area may be warranted.

## Acknowledgements

This study was supported by grants to BT from the Marsden Fund of New Zealand and the Canadian Institutes of Health Research [CIHR 365136].

## Data Availability

All clinical data are provided within the manuscript tables. Anonymized electrophysiological data are available from the authors upon request.

